# Evaluation of the cytotoxic potential of extracts from the genus *Passiflora* cultived in Brazil against cancer cells

**DOI:** 10.1101/337253

**Authors:** Ricardo Guimarães Amaral, Silvana Vieira Floresta Gomes, Ângelo Roberto Antoniolli, Maria Claudia dos Santos Luciano, Cláudia do Ó Pessoa, Luciana Nalone Andrade, Patrícia Severino, Geraldo Célio Brandão, Larissa Mendes Bomfim, Daniel Pereira Bezerra, Jorge Maurício David, Adriana Andrade Carvalho

## Abstract

This work aimed to evaluate the cytotoxic potential against cancer cells of *Passiflora* genus plant species cultivated in Brazil and identify the mechanism of cytotoxicity induced by the most promising extract. Leaf extracts from 14 *Passiflora* (P.) species were obtained ASE and *in vitro* cytotoxicity evaluated against cancer cell lines using MTT assay at a single concentration of 50 μg/mL. Additionally, the IC_50_ of the *P. alata* (ELPA) leaf extracts was determined against both tumor (HCT-116, SF-295, OVACAR-8, and HL-60), and non-tumor cells (PBMC). The ELPA flavonoids were identified by HPLC-DAD and UHPLC-MS/MS. The morphological analyses used light and fluorescence microscopy, and cell cycle and DNA fragmentation analyses used flow cytometry to determine the mechanism of cell death induced by ELPA in HL-60. Among the *Passiflora* leaf extracts evaluated; ELPA stood out with high cytotoxic activity, followed by *P. capsularis* and *P. quadrangulares* with varying high and low cytotoxic activity. ELPA presented high cytotoxic potency in HL-60 (IC_50_ 19.37 μg/mL), yet without cytotoxic activity against PBMC, suggesting selectivity for tumor cells. The cytotoxic activity of ELPA may well be linked to the presence of ten identified flavonoids. Cells treated with ELPA presented the hallmarks typical of apoptosis and necrosis, with cell cycle arrest in the G2/M phase. Conclusion: From among the studied species, ELPA presented greater cytotoxic activity, possibly a consequence of synergistic flavonoid action which induces cell death by apoptosis and necrosis.

## Introduction

Cancer is a progressive disease, characterized by a gradual accumulation of genetic mutations and epigenetic changes that cause cellular mechanism imbalances which progressively transform normal cells into malignant cells (1,2). Together with their associated treatment modalities for curative or palliative purposes, there are more than 100 types of cancers (3,4). Most therapeutic agents promote programmed cell death (apoptosis) in tumors. However, due to the relative similarity between malignant and normal cells, severe side effects are often observed; this is a limiting factor for therapeutics (2).

Despite existing treatment modalities, cancer is one of the most common causes of morbidity and mortality worldwide, with estimates that 14.1 million new cancers and 8.2 million cancer deaths occured in 2012, anticipating an increase of at least 70% by 2030, or 20 million new cases of cancer annually by 2025 (5,6).

Finding more effective and selective compounds that reduce this growing public health problem is a challenge, and nature is the alternative (7). Historically, natural plant and animal products have been the origin of most medicinal preparations and natural products continue to provide discovery clues towards pharmacologically active compounds, particularly anticancer agents (7,8). A continuing study beginning in 1981 through 2014 demonstrated that 83% of the world’s registered anticancer drugs were in one form or another either natural products, based therein, or mimetics (9). Despite these successes, most plant species have not been systematically investigated (8), and though Brazil has the greatest diversity of plant species in the world, discoverable tumor cell line cytotoxicities have been poorly studied (10).

A purpose of this study is to carry on the search for new natural product derived anticancer drugs. Within the gigantic diversity of species of the genus *Passiflora* (*P.*), some such as *P. edullis* Sims (11), *P. ligularis* Juss (12), *P. incarnata* (13–15), *P. molíssima* (16) and *P. tetandra* (17) have been described in the literature as cytotoxic to tumor cell lines, with *in vivo* chemopreventive or antitumor activity. This raises the hope that related species could exhibit similar activities.

Facing treatments with low curative power, a rising cancer mortality rate in the world, a great variety of plant species not yet having been studied in relation to their antitumor activity, and the fact that certain species of the genus *Passiflora* present anticancer activity, this study aimed to evaluate the cytotoxic potential of leaf extracts from fourteen plant species of the genus *Passiflora* against human cancer cell lines, and to identify the mechanism of cytotoxicity presented by most promising extract.

## Materials and Methods

### Collection of plant material

The leaves of fourteen species of *Passiflora* were collected in July 2011 in the Empresa Brasileira de Pesquisa Agropecuária (EMBRAPA), in the city of Cruz das Almas, Bahia, Brazil (12°04’10”S, 39°06’22”W). The species were identified, evaluated and registered in the active germoplasm bank (BGP) with access number BGP 08 (*P. giberti* N. E. Brown), BGP 32 (*P. maliformis* L.), BGP 77 (*P. cincinnata* Mast.), BGP 104 (*P. vitifolia* Kunth), BGP 105 (*P. tenuifila* Killip), BGP 107 (*P. morifolia* Mast.), BGP 109 (*P. galbana* Mast.), BGP 114 (*P. mucronata* Sessé & Moc.), BGP 125 (*P. capsularis* L.), BGP 152 (*P. suberosa* L.), BGP 157 (*P. quadrangularis* L.), BGP 163 (*P. alata* Curtis), BGP 170 (*P. malacophylla* Mast.), BGP 172 (*P. racemosa* Brot.) and BGP 237 (*P. setácea* DC.). The leaves of all species were air-dried at 30ºC for seven days in an airflow oven and then powdered; (only particles between 0.5 - 1.0 mm were utilized for the extractions).

### Preparation of extracts and samples

Extracts of the dried and powdered leaves were prepared by accelerated solvent extraction (ASE 100, Dionex Corporation, Sunnyvale, CA, USA). Each extract was prepared to weigh exactly 6.0 g *P. alata, P.capsularis, P. cincinnata, P. gibertii, P. maliformis, P. malacophylla, P. morifolia, P. mucronata, P. quadrangularis, P. racemosa, P. setacea, P. suberosa;* and 3.0 g of *P. vitifolia* and *P. tenuifila*. The plant material was extracted under optimized conditions in five extraction cycles with 64% (w/w) ethanol at 80°C, 1500 psi, and a static cycle timing of 10 min. The cells were then rinsed with fresh extraction solvent (100% of the extraction cell volume) and purged with N_2_ gas for 60 s. The solvent was removed under reduced pressure at 55°C to yield the corresponding crude extracts, which were stored under refrigeration and protected from light (18).

For HPLC analysis, hydroethanolic crude extract of *P. alata* was purified by solid-phase extraction (SPE) using the typical method with minor modifications (19). A C_18_ cartridge (Agilent SampliQ, 3 mL/200 mg) was conditioned with 3 mL of methanol, followed by 1 mL of water. Next, 2 mL of a methanolic solution of the sample (5 mg/mL) was added, and the flavonoid fraction was obtained by elution with 3mL of 60% (w/w) methanol. All samples were prepared and analyzed in triplicate.

### Identification of flavonoids in the extract of *P. alata*

#### HPLC-DAD

High-performance liquid chromatography (HPLC) *fingerprint* analysis was carried out using an LC system (Thermo Scientific Dionex Ultimate 3000; MA, USA) consisting of a Thermo Scientific Dionex Ultimate 3000 diode array detector (DAD), quaternary pump, on-line degasser and automatic sampler. The chromatographic separation of sample was achieved on a reversed-phase HPLC column (Waters XBridge™, BEH C_18_, 100 mm × 3.0 mm I.D., 2.5 μm particle size). For HPLC *fingerprint* chromatographic analysis, a binary gradient elution system composed of 0.2% (w/w) formic acid in water (solvent A) and acetonitrile (solvent B) was applied as follows: 0 - 30 min, 5 - 20% B. There re-quilibration duration between individual runs was 30 min. The injection volume was 10 µL per sample, the flow rate was 0.6 mL/min and the column temperature was maintained at 30 ºC. The flavonoids were detected at 337 nm, and the UV spectra of individual peaks were recorded within a range of 190–400nm. Data were acquired and processed with Chromeleon *software*. The main flavonoids were identified in the extract of *P. alata* based on their retention times (t_R_), coinjection of the samples with standards and comparison of their UV adsorption spectra (18).

#### UHPLC-MS/MS

Ultra-high performance liquid chromatography tandem mass spectrometry (UHPLC-MS/MS) was utilized to identify and confirm chromatogram peaks, which was comprised of an ACQUITYTM UHPLC system (Waters Corp., Milford, MA, USA). Separation was performed with an ACQUITY BEH C_18_ column (50 mm × 2 mm, 1.7 µm; Waters, USA). The mobile phase was a mixture of 0.1% formic acid (A) and acetonitrile in 0.1% formic acid (B), with an linear gradient elution as follows: 0–11 min, 5–95% B. The injection volume was 4 µL. The flow rate was set at 0.30 mL/min. The UV spectra were registered from 190 to 450 nm. Eluted compounds were detected from *m/z* 100 to 1000 using a Waters ACQUITY^®^ TQD tandem quadrupole mass spectrometer equipped with an electrospray ionization (ESI) source in the negative mode using the following instrument settings: capillary voltage 3500V; capillary temperature 320°C; source voltage 5 kV; vaporizer temperature 320°C; corona needle current 5 mA; and sheath gas nitrogen 27 psi. Analyses were run in the full scan mode (100– 2000 Da). Ion spray voltage: −4 kV; orifice voltage: −60 V. The identification of flavonoids was based on the comparison of the molecular formula with that of the published data, further confirmation was performed by illuminating the quasi-molecular ions and key flavonoids fragmentations, in particular for those isomeric.

### Human cancer cell lines and non-tumor cells

The cytotoxicity of these *Passiflora* species extracts was tested against colon carcinoma (HCT-116), glioblastoma (SF-295), ovarian adenocarcinoma (OVCAR-8) and pro-myelocytic leukemia (HL-60) human cancer cell lines, all obtained from the National Cancer Institute, Bethesda, MD, USA. The cells were grown in RPMI-1640 medium supplemented with 10% fetal bovine serum, two mM glutamine, 100 µg/mL streptomycin and 100 U/mL penicillin incubated at 37°C in a 5% CO_2_ atmosphere. Peripheral blood mononuclear cells (PBMCs) were isolated from a sample of about 3 mL of human blood plus 5 mL of saline. The steps up to isolation included the addition of 3 mL of Ficoll followed by 15 minutes of centrifugation at 1500 rpm, and aspiration of the PBMCs present in the intermediate region between the red blood cells and the plasma. The PBMC suspension was transferred to another tube which was added with saline to the 11 mL volume and centrifuged for 5 minutes at 1000 rpm. The supernatant was discarded, and the PBMC pellet was re-suspended in complete medium (RPMI 1640 plus 20% fetal bovine serum and 10 μg/mL ConA) and counted in a Neubauer chamber for further dilution and plating.

### Determination of the cytotoxic effect of Passiflora extracts on cancer cell lines

The MTT assay was used to determine the cytotoxic effect of the 14 extracts of Passiflora leaves against tumor cell lines (20). For all experiments, HCT-116 cells (0,7 × 10^5^ cells/mL in 100 µL medium), SF-295 and OVCAR-8 cells (0.1 × 10^6^ cells/mL in 100 µL medium) were seeded in 96-well plates and incubated in a humidified chamber with 5% CO_2_ at 37°C for 24 hours. After this the 14 extracts of Passiflora were solubilized in dimethyl sulfoxide (DMSO, 0.7%), and added to each well at the concentration of 50 µg/mL. The cells were then incubated for 72 h at 37°C in a 5% CO_2_ atmosphere. After 72 hours, MTT (0.5 mg/mL) was added, followed by incubation at 37°C in an atmosphere of 5% CO_2_ for 3 hours. After incubation, the formazan product was dissolved in 150 μL DMSO, and absorbance was measured using a multi-plate reader (DTX 880 Multimode Detector, Beckman Coulter Inc.). The treatment’s effects were expressed as the percentage of control absorbance of reduced dye at 595 nm. All absorbance values were converted into a cell growth inhibition percentage (GI %) by the following formula:
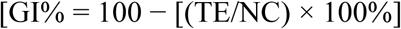
In which, NC is the absorbance for the negative control, and TE is the absorbance in the presence of the tested extracts.

The median inhibitory concentration able to induce 50% of maximal effect (IC_50_) was determined for the extract that caused the greatest GI% against HCT-116, SF-295 and OVCAR-8 The same protocol for the same cells, adding the cell line HL-60 (0.3 × 10^6^ cells/mL in 100 µL medium) was used to determine the IC_50_, varying only the concentration of the compound from 0 to 50 µg/mL, and the potency of the extract. The experiment was performed as three independent experiments, and doxorubicin (0.3 μg/mL) was used as a positive control (Doxorubicin, purity > 98%; Sigma Chemical Co., St. Louis, MO, USA).

### Determination of the effect of *P. alata* leaf extract in non-tumor cells

The alamar blue assay was used to determine the effect of extract from the leaves of *P. alata* (ELPA) against proliferation of non-tumor cells (PBMCs) obtained from peripheral blood from healthy human volunteers (21). Initially, the cells were plated in 96-well plates (100 μL/well of a solution of 3 × 10^6^ cells/mL). After 24 hours of incubation, extract from the leaves of *P. alata* (0.39 to 50 µg/mL) was dissolved in 0.3% DMSO, added to each well and incubated for 72 hours. Doxorubicin (0.03 - 0.25 μg/mL) was used as a positive control and 0.3% DMSO was used as a negative control. Four hours before the end of the incubation period, 20 μL of alamar blue stock solution (0.312 mg/mL) (resazurin) was added to each well. Absorbances were measured at wavelengths of 570 nm (reduced) and 595 nm (oxidized) using a plate reader (DTX 880 Multimode Detector, Beckman Coulter™).

### Determination of the cytotoxicity mechanism induced by *P. alata* leaf extract

#### Morphological Analysis by light microscopy

Differential staining with hematoxylin/eosin was used for morphological analysis. The HL-60 cells, plated at 0.3 × 10^6^ cells/mL were incubated for 72 h with *P. alata* leaf extract (9.69, 19.77, and 38.74 μg/mL) and examined under an inversion microscope. To observe the morphology, 50 μL of the cell suspension was added to the slide centrifuge (Shandon Southern Cytospin™). After cell adhesion on the slide, fixation with methanol for 1 minute was completed, hematoxylin followed by eosin was used. Doxorubicin (0.3 μg/mL) was used as a positive control (22).

#### Morphological analysis by fluorescence microscopy

Acridine orange/ethidium bromide (AO/EB) (Sigma Aldrich) staining in HL-60 cells was performed to evaluate the pattern of cell death induced by *P. alata* leaf extract. Cells of the HL-60 strain were plated at 0.3 × 10^6^ cells/mL, and then incubated for 72 h with the extract (9.69, 19.77, and 38.74 μg/mL). The cell suspension was transferred to an Eppendorf tube and centrifuged for 5 min at low speed (500 rpm). The supernatant was discarded and the cells re-suspended in 20 μL of PBS solution. Then 1 μL of the ethidium bromide/acridine orange solution was added to each tube and placed under a fluorescence microscope to observe the cellular events. Doxorubicin 0.3 μg/mL was used as a positive control (23).

#### Analysis of cell cycle and DNA fragmentation

Cell cycle analysis was performed to determine cell content, which reflects the cell cycle phases (G0/G1, S, and G2/M). For this, HL-60 cells were plated in 24-well plates at the concentration of 0.3 × 10^6^ cells/mL and incubated for 72 h with *P. alata* leaf extract (9.69, 19.77 and 38.74 μg/mL). Doxorubicin (0.3 μg/mL) was used as a positive control. After the incubation period an aliquot of the cell suspension (50 μL) was transferred to an Eppendorff tube and incubated for another 30 minutes with 100 μL of a lysis solution containing propidium iodide (50 μg/mL), Triton X-100 (0.1%) and sodium citrate (0.1%), in the absence of light and at 37°C (24). The samples were then analyzed in the program Guava Express Plus, where the number of cells in each phase of the cell cycle was obtained along with the number of cells presenting fragmented DNA.

#### Statistical Analysis

Data are presented with mean ± standard error of the mean (SEM), standard deviation (SD), or IC_50_ values; confidence intervals (CI 95%) were obtained by nonlinear regression. Differences among the experimental groups were compared by one-way variance analysis (ANOVA), followed by Newman-Keuls test (*p* < 0.05). All analyses were carried out using the Graphpad program (Intuitive Software for Science, San Diego, CA, USA).

## Results and Discussion

Based on the fact that some species of the genus *Passiflora* have cytotoxic activity against tumor cell lines and that relatively few species have been evaluated for their possible anticancer activity, we selected 14 species of *Passiflora* cultivated in Brazil, produced by ASE method to evaluate their potential *in vitro* cytotoxic activity.

The cell growth inhibition percentage (GI%) of the *Passiflora* extracts was evaluated against three human cancer cell lines (HCT-116, OVACAR-8, and SF-295) in a single concentration of 50 µg/mL and measured by MTT assay after 72 hours of incubation (Table 1). The results were analyzed for each cell line tested using a GI% scale as follows: samples with low cytotoxic activity have GI < 50%; samples with moderate cytotoxic activity have GI between 50% and 75%, and samples with high cytotoxic activity have GI > 75% (25). The results showed that the extracts of *P. cincinnata* (3), *P. gibertii* (4), *P. maliforms* (5), *P. mallacophyla* (6), *P. murchronata* (7), *P. morifolia* (8) *P. racemosa* (10), *P. setacea* (11), *P. suberosa* (12), *P. tenuifila* (13) and *P. vitifolia* (14) have low or nonexistent cytotoxic activity in all human cancer cell lines tested. Extracts from *P. capsularis* (2) and *P. quadrangulares* (9) have high activity against HCT-116, low activity against SF-295, and cytotoxic activity from low (*P. quadrangulares*) to moderate (*P. capsularis*) against OVACAR-8. The extract from the leaves of *P. alata* (1) presented high cytotoxic activity in all human cancer cell lines analyzed. Therefore, among the *Passiflora* leaf extracts evaluated *P. alata* (1) stands out with promising cytotoxic activity, followed by *P. capsularis* (2) and *P. quadrangulares* (9).

**Table 1.**
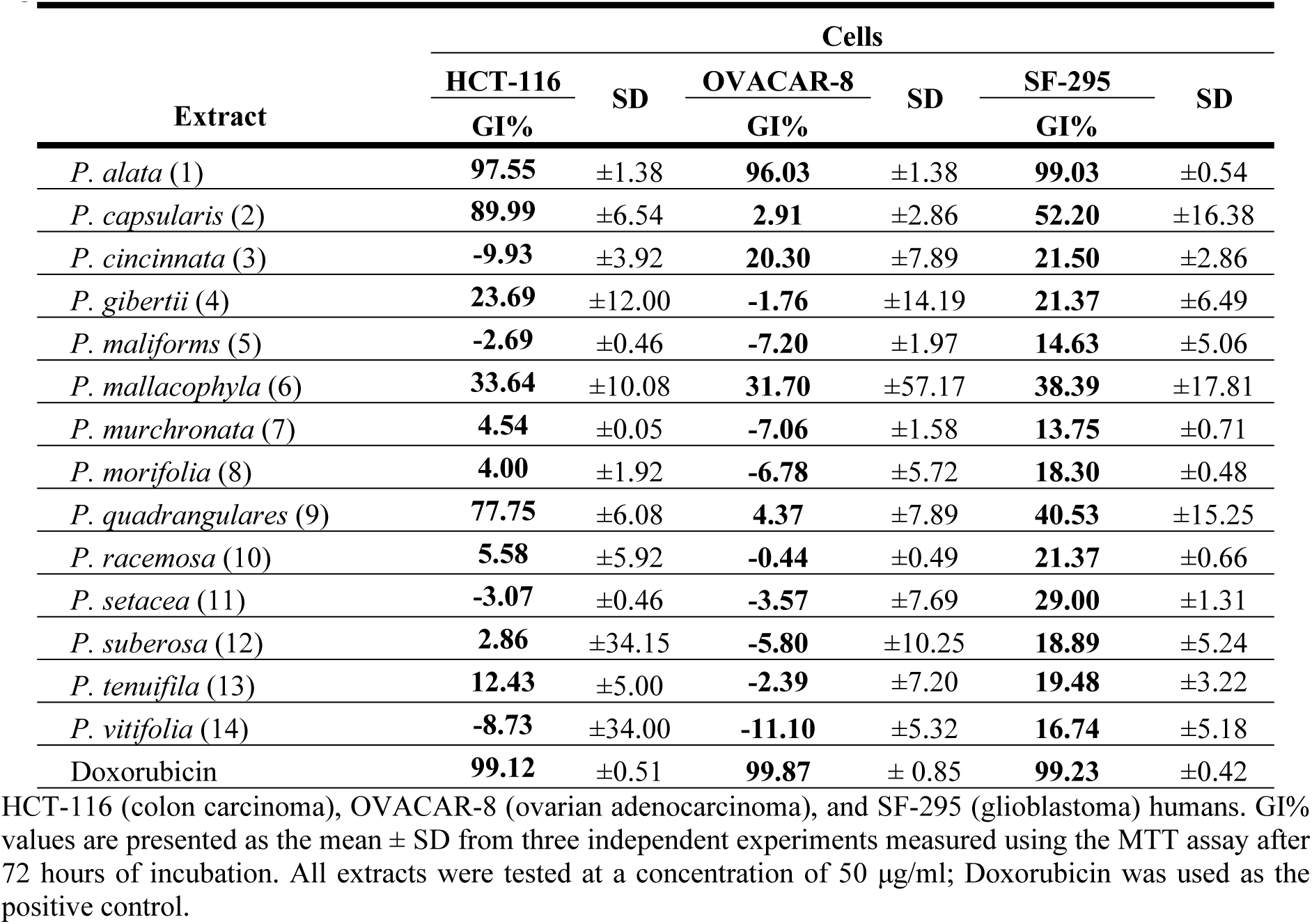
Cell growth inhibition percentage (GI%) of leaf extracts from 14 species of *Passiflora* against human tumor cell lines.

| Extract | Cells |  |  |  |  |  |
| --- | --- | --- | --- | --- | --- | --- |
|  | HCT-116 | SD | OVACAR-8 | SD | SF-295 | SD |
|  | GI% |  | GI% |  | GI% |  |
| *P. alata* (1) | **97.55** | ±1.38 | **96.03** | ±1.38 | **99.03** | ±0.54 |
| *P. capsularis* (2) | **89.99** | ±6.54 | **2.91** | ±2.86 | **52.20** | ±16.38 |
| *P. cincinnata* (3) | **-9.93** | ±3.92 | **20.30** | ±7.89 | **21.50** | ±2.86 |
| *P. gibertii* (4) | **23.69** | ±12.00 | **-1.76** | ±14.19 | **21.37** | ±6.49 |
| *P. maliformis* (5) | **-2.69** | ±0.46 | **-7.20** | ±1.97 | **14.63** | ±5.06 |
| *P. mallacophylla* (6) | **33.64** | ±10.08 | **31.70** | ±57.17 | **38.39** | ±17.81 |
| *P. murchronata* (7) | **4.54** | ±0.05 | **-7.06** | ±1.58 | **13.75** | ±0.71 |
| *P. morifolia* (8) | **4.00** | ±1.92 | **-6.78** | ±5.72 | **18.30** | ±0.48 |
| *P. quadrangulares* (9) | **77.75** | ±6.08 | **4.37** | ±7.89 | **40.53** | ±15.25 |
| *P. racemosa* (10) | **5.58** | ±5.92 | **-0.44** | ±0.49 | **21.37** | ±0.66 |
| *P. setacea* (11) | **-3.07** | ±0.46 | **-3.57** | ±7.69 | **29.00** | ±1.31 |
| *P. suberosa* (12) | **2.86** | ±34.15 | **-5.80** | ±10.25 | **18.89** | ±5.24 |
| *P. tenuifila* (13) | **12.43** | ±5.00 | **-2.39** | ±7.20 | **19.48** | ±3.22 |
| *P. vitifolia* (14) | **-8.73** | ±34.00 | **-11.10** | ±5.32 | **16.74** | ±5.18 |
| Doxorubicin | **99.12** | ±0.51 | **99.87** | ± 0.85 | **99.23** | ±0.42 |

HCT-116 (colon carcinoma), OVACAR-8 (ovarian adenocarcinoma), and SF-295 (glioblastoma) humans. GI% values are presented as the mean ± SD from three independent experiments measured using the MTT assay after 72 hours of incubation. All extracts were tested at a concentration of 50 µg/ml; Doxorubicin was used as the positive control.

Although none of the three species highlighted in the present study had been previously evaluated for their cytotoxic activity against cancer cells, other species of the genus *Passiflora* have been. The cytotoxic or antiproliferative activity of species of the genus *Passiflora* was evidenced for the first time by Perry et al. (1991) (17) through study of *P. tetandra* leaf extract, demonstrating cytotoxic activity against murine leukemic cell line (P388), being possibly mediated by the extract component 4-hydroxy-2-cyclopentane. The ethanolic extract of *P. ligularis* Juss. and the methanolic extract of the *P. edullis* fruit have been respectively described as presenting activity against human hepatocellular carcinoma (Hep3B) and leukemic cell line (CCRF-CEM) (11,12).

Based on the promising cytotoxic activity observed in the *P. alata* (ELPA) leaf extract we determined the median inhibitory concentration able to induce 50% of maximal effect (IC_50_) against four human cancer cell lines (HCT-116, HL-60, OVCAR-8, and SF-295) through the MTT method. The results demonstrated that ELPA has cytotoxic activity in the four cancer cell lines evaluated, respectively being most potent for the cell line HL-60 (IC_50_ of 19.37 µg/mL), followed by HCT-116 (IC_50_ of 20.79 µg/mL), SF-295 (IC_50_ of 21.87 µg/mL), and OVCAR-8 (IC_50_ of 28.26 µg/mL) (Table 2). According to the preclinical cytotoxic drug screening program based on the US National Cancer Institute program, only extracts with IC_50_ values below 30 μg/mL in assays with tumor cell lines are considered promising for the development of anticancer drugs (25,26). Thus, ELPA presents promising *in vitro* cytotoxic activity for further study.

**Table 2.** Evaluation of the cytotoxic activity of *P. alata* leaf extract (ELPA) against tumor and non-tumor cells.

| Cells | Histotype | ELPA<br>µg/mL | Doxorubicin<br>µg/mL |
| --- | --- | --- | --- |
| **HL-60** | Promyelocytic leukemia | 19.37 | 0.21 |
|  |  | 15.46 – 24.27 | 0.14 – 0.31 |
| **HCT-116** | Colon carcinoma | 20.79 | 0.02 |
|  |  | 19.06 – 22.68 | 0.01 – 0.04 |
| **SF-295** | Glioblastoma | 21.87 | 0.24 |
|  |  | 18.83 – 25.41 | 0.17 – 0.36 |
| **OVCAR-8** | Ovarian adenocarcinoma | 28.26 | 1.10 |
|  |  | 22.13 – 29.40 | 1.00 – 1.50 |
| **PBMC** | Peripheral blood mononuclear cells | > 50 | 0.77 |
|  |  |  | 0.43 – 1.40 |

Values were expressed as IC<sub>50</sub> and obtained by nonlinear regression from three independent experiments, performed in triplicate and measured with the MTT assay after 72 h of incubation with the ELPA; values are means with 95% confidence limits in parentheses. Doxorubicin was used as positive control.

For a natural compound to progress as a potential anticancer drug candidate, it is necessary to determine the degree of specificity of the drug by evaluating *in vitro* cytotoxic activity against non-tumor cells (26,27). For this, ELPA was subjected to *in vitro* cytotoxic activity evaluation against non-tumor cells (PBMC); the results demonstrated that the compound shows no cytotoxic activity against PBMC (Table 2).

The cytotoxic activity of natural bioactive products against cancer cells is attributed to the chemical composition of the product, and this practice of identification is the most successful source of potential drug discovery and development (7).

A total of 4 flavonoids were identified by HPLC-DAD, and 8 flavonoids were identified by UHPLC-MS/MS in the negative mode. They are summarized along with their retention time, molecular formula and MS/MS fragments as shown in Table 3. HPLC fingerprint chromatogram of ELPA with 4 identified compounds are show in Figure 1.

**Figure 1.**
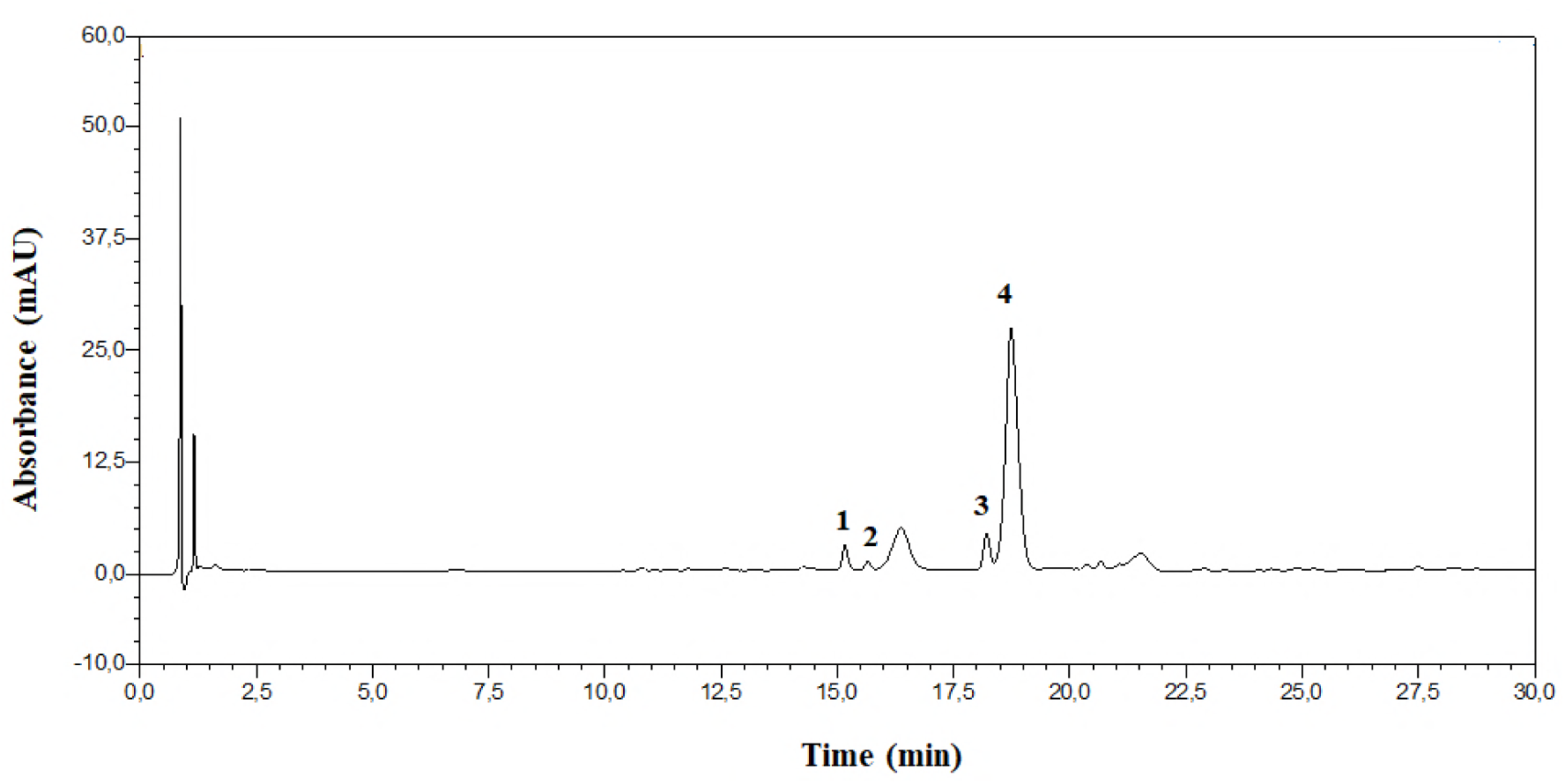
HPLC fingerprint chromatogram (λ=337 nm) of ELPA. Mobile phase: ACN (B):0.2% (w/w) formic acid (A) (0–30 min, 5–20% of B). Flow rate: 0.6 mL/min. Injection volume: 0.6 μL. Column temperature: 30 °C. Peaks: 1 — isoorientin; 2 — orientin; 3 — vitexin; 4 — isovitexin.

**Table 3.** Identification of the chemical constituents from ELPA by UHPLC-MS/MS.

| Peak No. | Retention time (min) | $\lambda_{\text{max}}$ (nm) | [M-H]<sup>-</sup> m/z | MS/MS fragments ions | Identification |
| --- | --- | --- | --- | --- | --- |
| 1 | 2.65 | 269; 338 | 431.43 | 340.9; 311.0; 295.1; 282.8 | Vitexin |
| 2 | 2.76 | 270; 339 | 431.12 | 341.2; 311.1; 295.0; 282.9 | Isovitexin |
| 3 | 2.37 | 266; 345 | 446.72 | 356.8; 327.2; 297.0; 255.0 | Orientin |
| 4 | 2.48 | 268; 338 | 446.72 | 356.8; 327.2; 298.9; 284.8 | Isoorientin |
| 5 | 2.80 | 270; 343 | 461.37 | 341.0; 313.3; 297.8 | 8-C-Glucosyldiosmetin |
| 6 | 2.89 | 270; 343 | 461.05 | 341.2; 298.0 | 6-C-Glucosyldiosmetin |
| 7 | 2.71 | 269; 339 | 607.46 | 487.0; 432.3; 340.7; 322.9; 298.0 | 6-C-Glucoronilvitexin |
| 8 | 2.78 | 269; 340 | 607.12 | 487.0; 432.3; 341.4; 322.9; 298.0 | 8-C-Glucoronilisoovitexin |

Flavonoid from *Passiflora* spp. are usually *C*-glycosylated with one or more sugar units. Different fragmentation patterns were observed in MS/MS experiments for flavone *C*-glycosides. Based on MS and UV/Vis spectra, all identified flavones were vitexin, isovitexin, orientin, isorientin, 8-*C*-glucosyldiosmetin, 6-*C*-glucosyldiosmetin, 6-*C*-glucoronylvitexin and 8-*C*-glucoronylisovitexin.

Thus, compounds 1 and 2 (t_R_ = 2.65 and 2.76 min) were identified as vitexin and isovitexin, once the [M-H]-pseudo molecular ions were registerated at *m/z* 431 and the fragmentation pattern, compared with the literature are also in accordance with these two compounds. In the same way compounds 3 and 4 (t_R_ = 2.37 and 2.48 min) were identified as orientin and isoorientin, also due the pseudo [M-H]-ions at *m/z* 446. These compounds (1–4) indicated similarities of fragmentation patterns due to dehydration and cross-ring cleavages of the glucose moiety, i.e., 0.2 cross-ring cleavage (− 120 amu) and 0.3 cross-ring cleavage (− 90 amu) (Figure 2) (28).

**Figure 2.**
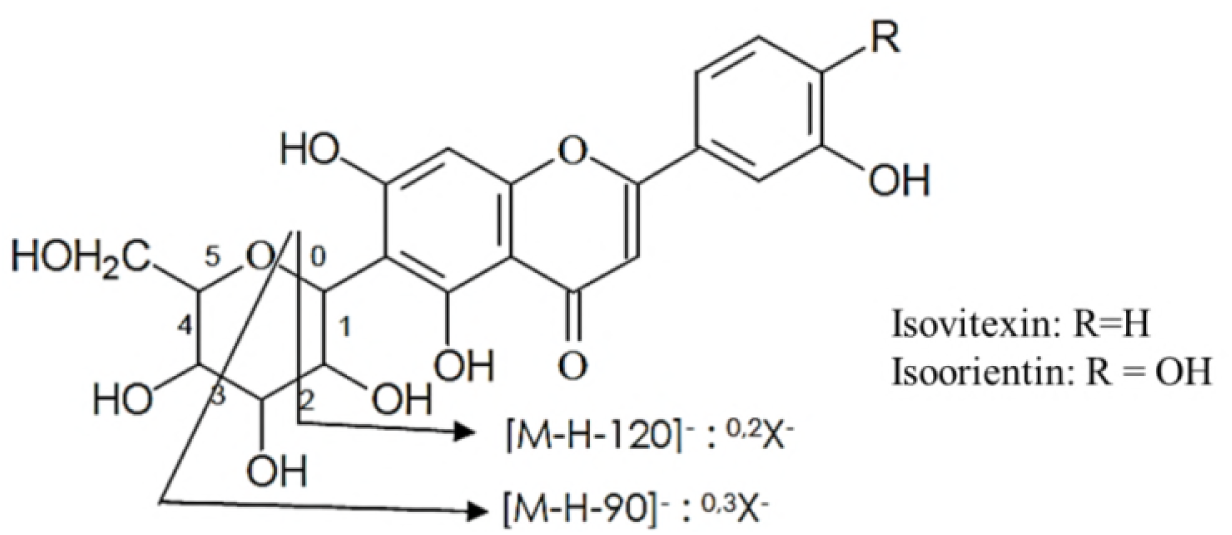
Fragmentation of isovitexin and isoorientin.

Compounds 5 and 6 (t_R_ = 2.80 and 2.89 min) were identified as 8-*C*-glucosyldiosmetin and 6-*C*-glucosyldiosmetin, [M-H]-ions presented at *m/z* 461. In the MS/MS spectra the fragments were at *m/z* 341 ^0,2^X-[M-H-120]-, which was attributed to the loss of a glucose moiety.

Compounds 7 and 8 (t_R_ = 2.71 and 2.78 min) exhibited [M-H]-ions at *m/z* 607. They were identified as 6-*C*-Glucoronylvitexin and 8-*C*-Glucoronylisovitexin. In the MS/MS spectrum, the same fragment ion at *m/z* 487 ^0,2^X-[M-H-120]-indicated the loss of a glucose moiety. The fragment ion at *m/z* 432 suggesting the loss of a glucuronic acid residue [M-H-175]- (29–31).

The extractive process used for the ELPA was ASE, developed to maximize the yield of the extract and simultaneously the content of flavonoids (18). According Pearson et al. (2010) (32) and Saha et al. (2015) (33), the process is promising, because it offers advantages over other extraction processes such as: easy automation, faster sample analysis, better reproducibility, less solvent required, and sample maintenance in an environment without light and free of oxygen. Accelerated extraction systems also allow the operator to control the temperature, pressure, extraction time and number of extractions, which may increase the number of compounds extracted from the plant when optimized. Thus, ASE could have influenced the optimization of extraction and increased the number of flavonoids extracted from ELPA.

The observed cytotoxic activity of ELPA demonstrated in this study might be attributed to the mixture of various flavonoids found in the extract. Flavonoids have the potential to modify many biological cancer events, such as apoptosis, vascularization, cell differentiation and proliferation (34). Flavonoids found in ELPA such as vitexin and isovitexin also exert chemotherapeutic potential against breast, hepatic, colorectal, lung, skin, oral, prostate, cervix, ovary, esophagus and leukemic cancers (35). Orientin shows an inhibitory effect on the proliferation of esophageal cancer cells (EC-109) and isoorientin demonstrated the ability to inhibit proliferation of liver cancer cells (HepG2) (36–38). Diosmetin, found in ELPA conjugated with C-glucosyl groups (8-C-Glucosyldiosmetin and 6-C-Glucosyldiosmetin), is described as having *in vitro* anticancer activity against several tumor cells lines, with potent inhibition against leukemic cells (P-388), and lesser cytotoxic activity against HepG2, Hep3B, MDA-MB-231, MCF-7, A549, and Ca9-22 tumor cells (39). These data may justify the *in vitro* cytotoxic effect presented by ELPA.

For drug discovery, cell death related testing is paramount in oncology because resistance to dying is a hallmark of cancer cells (40). Knowing the cytotoxic activity demonstrated by ELPA, we decided to identify the mechanism of the cytotoxicity induced by the extract. For this, HL-60 neoplastic cells were selected for subsequent testing based on the fact that the lineage is among the most widely used cellular models of myeloid origin, and simultaneously was the lineage against which ELPA demonstrated its highest cytotoxic potency. Three concentrations of ELPA, ½ IC_50_ (9.69 μg/mL), IC_50_ (19.37 μg/mL) and 2 × IC_50_ (38.74 μg/mL) were chosen.

To ascertain the cellular death process in cancer cells, two tests were performed, hematoxylin-eosin coloration analyzed by light microscopy, and ethidium bromide/acridine orange coloration using fluorescence microscopy. After 72 h incubation, the effect of ELPA was evaluated based on cell morphology using hematoxylin-eosin. The HL-60 cells treated with ELPA showed the characteristic morphology of cell death induced by apoptosis (reduction in cell volume, chromatin condensation, and nuclear fragmentation), and necrosis (membrane disruption and cell swelling) as shown in Figure 3.

**Figure 3.**
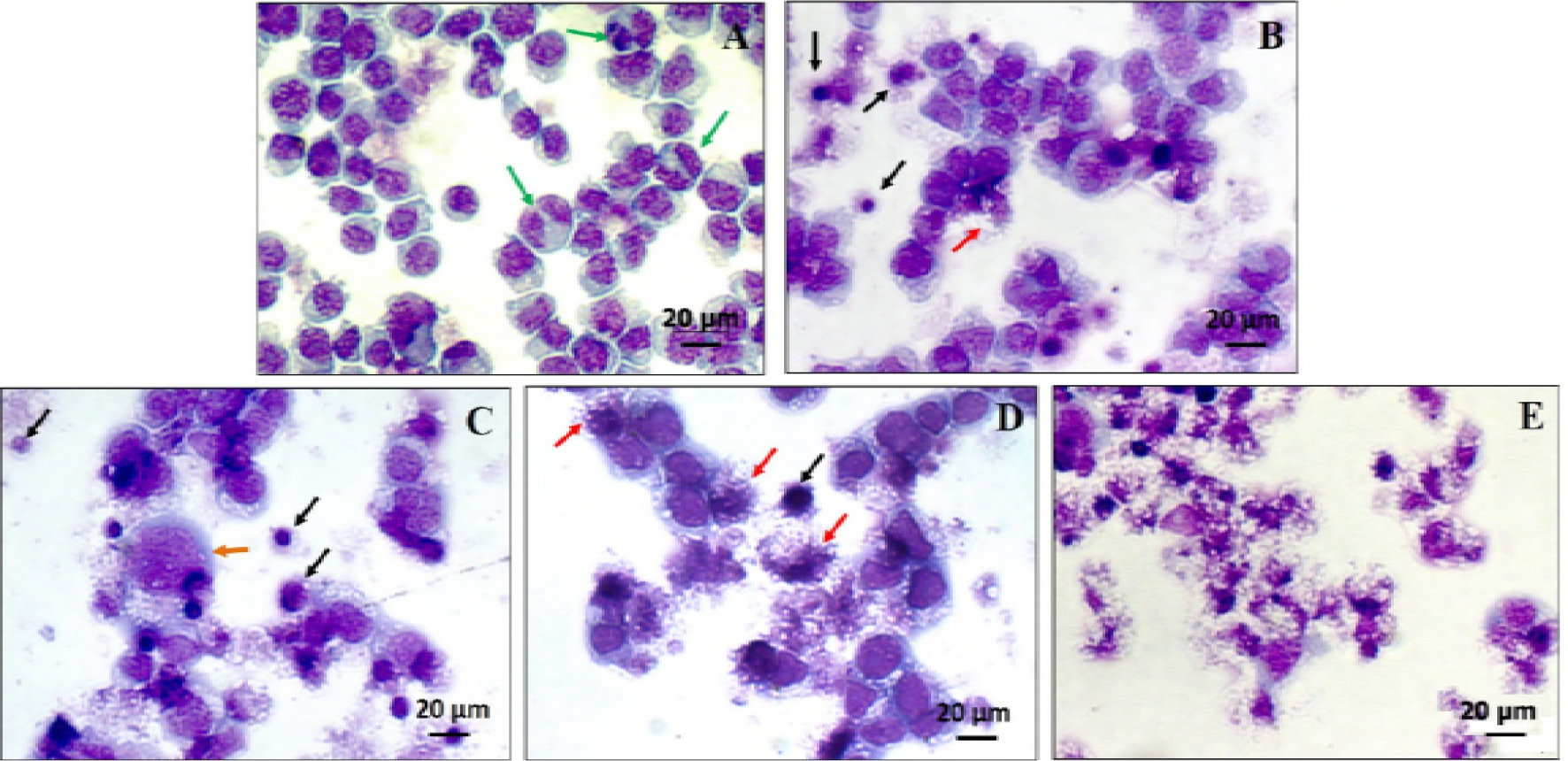
Photomicrography of human tumor cells (HL-60) submitted to hematoxylin/eosin differential staining after 72 hours of incubation and analyzed under a light microscope (100X). **A** - negative control, **B** - cells treated with ELPA 9.69 µg/mL, **C** - cells treated with ELPA 19.77 µg/mL, **D** - cells treated with ELPA 38,74 µg/mL and **E** - cells treated with doxorubicin. Black arrows indicate cell volume reduction and nuclear fragmentation; red arrows indicate loss of membrane integrity, green arrows indicate cells in mitosis and orange arrow cell swelling.

To confirm the light microscopy findings, we performed a morphological analysis of cells treated with ELPA by staining and examining cells with AO/EB by fluorescence microscopy. The percentages of viable, apoptotic and necrotic cells were calculated. After 72 h, HL-60 cells treated with ELPA showed a reduction in the percentage of viable cells for the concentrations of 9.69 µg/mL (48.82 ± 3.76%), 19.77 µg/mL (41.88 ± 2.58%) and 38.74 µg/mL (34.74 ± 1.25%) when compared to the NC group (91.67 ± 1.72%). The percentage of cells in apoptosis increased for the concentrations of 9.69 µg/mL (30.74 ± 3.22%), 19.77 µg/mL (32.44 ± 3.23%) and 38.74 µg/mL (35.88 ± 3.06%) when compared to the NC group (4.95 ± 0.98%). Another process of cell death observed was the increase in the percentage of cells in necrosis, for the concentrations of 9.69 µg/mL (20.44 ± 5.86%), 19.77 µg/mL (26.88 ± 5.03%) and 38.74 µg/mL (29.45 ± 3.92%) when compared to the NC group (3.17 ± 1.18%) as shown in Figure 4.

**Figure 4.**
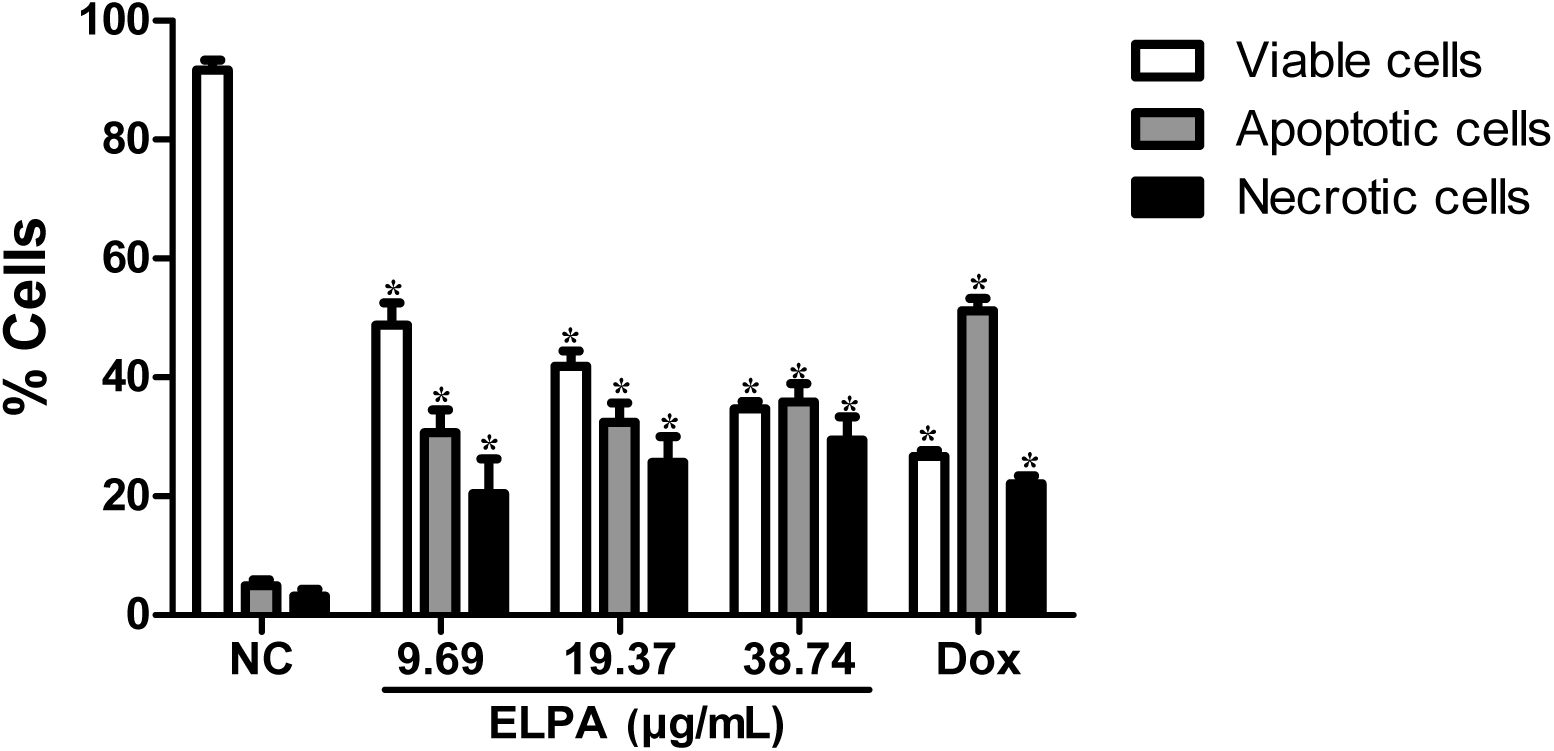
Identification of the type of cell death induced by *P. alata* leaf extract (EFPA) against HL-60. Viable cells (white bar), apoptosis (gray bar) and necrosis (black bar) were determined by fluorescence microscopy using ethidium bromide/acridine orange after 72 hours of incubation. The negative control (NC) was treated with DMSO 0.3%, and doxorubicin (Dox), used as a positive control. Data are presented as mean ± SEM of three independent experiments evaluated by one-way variance analysis (ANOVA) with a *Student Newman Keuls* post-test. *p < 0.05 compared to the NC group, and #p < 0.05 compared to the Dox group.

These data reaffirm the results found in the morphological analyses using light microscopy; demonstrating that ELPA can cause cell death by apoptotic and necrotic processes. Apoptosis is a programmed cell death process, and many studies suggest the activation of apoptosis as one of the most effective forms of chemotherapy to prevent the development and progression of cancer (41,42). Photodynamic treatment and certain antineoplastic agents, such as β-lapachone and DNA alkylating agents induce cell death by apoptosis and simultaneously induce cell death by necrosis in a variety of cancer cells (43,44). Considering the two cell death processes involved, ELPA is a promising candidate for *in vivo* evaluation in animal experimental tumor models.

The morphological analysis in HL-60 indicated that apoptosis is one of the mechanisms of death involved in the cytotoxicity observed in ELPA, we resolved to determine DNA fragmentation and alteration in the cell cycle through flow cytometry. The results concerning cell cycle progression showed that the ELPA at all concentrations tested showed a significant increase in the percentage of cells in the G2/M phase, being indicative of cell cycle arrest in the specific phase (Table 4). Assessing DNA fragmentation (SubG1), the data demonstrated a significant (p < 0.05) increase in fragmentation in the ELPA-treated groups at concentrations of 19.97 and 38.74 μg/mL (Table 4). Thus, cell cycle arrest may be related to ELPA-promoted DNA damage and subsequent DNA repair attempts. The lesion appears to be intense, as the DNA repair does not occur, and apoptosis is triggered.

**Table 4.** Effect of *P. alata* leaf extract (ELPA) on nuclear DNA content in HL-60 cells determined by flow cytometry.

| Treatment | Concentration<br>$\mu\text{g/mL}$ | DNA Content (%) | | | |
| --- | --- | --- | --- | --- | --- |
|  |  | Sub-G1 | G<sub>1</sub> | S | G<sub>2</sub>/M |
| NC | - | 12.92 ± 1.76 | 40.13 ± 1.32 | 36.96 ± 3.46 | 8.66 ± 3.85 |
| ELPA | 9.69 | 17.79 ± 1.82 | 34.56 ± 6.15 | 31.19 ± 3.40 | 16.87 ± 1.12* |
| ELPA | 19.77 | 25.08 ± 3.13* | 25.03 ± 1.83* | 26.51 ± 2.12* | 23.77 ± 0.73* |
| ELPA | 38.74 | 40.05 ± 4.14* | 16.69 ± 2.28* | 19.08 ± 1.34* | 24.21 ± 4.69* |
| Doxorubicin | 0.3 | 55.42 ± 4.59* | 5.99 ± 0.99* | 10.59 ± 1.80* | 26.38 ± 0.41* |

The data presented correspond to the mean ± standard error of the mean of two independent experiments performed in duplicate and evaluated by one-way analysis of variance with *Student Newman Keuls* post-test. \* $p < 0.05$ compared to the negative control.

## Conclusions

In conclusion, we evaluated the cytotoxic potential of extracts from 14 *Passiflora* species obtained by ASE. ELPA presented greater cytotoxic potential against tumor cells, without cytotoxicity in non-tumor human cells. The *in vitro* cytotoxic activity may be a consequence of synergistic action between the flavonoids found in ELPA, promoting cell cycle arrest in the G2/M phase and inducing cell death by apoptosis and necrosis. This work thus demonstrates the potential of Brazilian plants and opens perspectives for ELPA evaluation in experimental *in vivo* tumor models.

## Acknowledgments

This research was supported by the Coordenação de Aperfeiçoamento de Pessoal de Nível Superior (CAPES), and the Fundação de Apoio à Pesquisa e Inovação Tecnológica do Estado de Sergipe (FAPITEC/SE).

## References

1. You JS, Jones PA. Review Cancer Genetics and Epigenetics : Two Sides of the Same Coin ? Cancer Cell. 2012;22(1):9–20.

2. Hedvat M, Emdad L. K. Das S, Kim K, Dasgupta S, Thomas S, et al. Selected Approaches for Rational Drug Design and High Throughput Screening to Identify Anti-Cancer Molecules. Anticancer Agents Med Chem. 2012;12(9):1143–55.

3. Ge C, Li R, Song X, Qin S. Advances in evidence-based cancer adoptive cell therapy. Chinese Clin Oncol. 2017;6(2):18–18.

4. Palumbo MO, Kavan P, Miller WH, Panasci L, Assouline S, Johnson N, et al. Systemic cancer therapy: Achievements and challenges that lie ahead. Front Pharmacol. 2013;4 MAY(May):1–9.

5. Antoni S, Soerjomataram I, Møller B, Bray F, Ferlay J. An assessment of GLOBOCAN methods for deriving national estimates of cancer incidence. Bull World Health Organ. 2016;94(3):174–84.

6. Ferlay J, Soerjomataram I, Dikshit R, Eser S, Mathers C, Rebelo M, et al. Cancer incidence and mortality worldwide: Sources, methods and major patterns in GLOBOCAN 2012. Int J Cancer. 2015;136(5):E359–86.

7. Dias DA, Urban S, Roessner U. A Historical Overview of Natural Products in Drug Discovery. Metabolites. 2012;2(4):303–36.

8. Harvey AL, Edrada-Ebel R, Quinn RJ. The re-emergence of natural products for drug discovery in the genomics era. Nat Rev Drug Discov. 2015;14(2):111–29.

9. Newman DJ, Cragg GM. Natural Products as Sources of New Drugs from 1981 to 2014. J Nat Prod. 2016;79(3):629–61.

10. Ferreira PMP, Farias DF, Viana MP, Souza TM, Vasconcelos IM, Soares BM, et al. Study of the antiproliferative potential of seed extracts from Northeastern Brazilian plants. An Acad Bras Cienc. 2011;83(3):1045–58.

11. Kuete V, Dzotam JK, Voukeng IK, Fankam AG, Efferth T. Cytotoxicity of methanol extracts of Annona muricata, Passiflora edulis and nine other Cameroonian medicinal plants towards multi-factorial drug-resistant cancer cell lines. Springerplus. 2016;5(1):1666.

12. Carraz M, Lavergne C, Jullian V, Wright M, Gairin JE, Gonzales De La Cruz M, et al. Antiproliferative activity and phenotypic modification induced by selected Peruvian medicinal plants on human hepatocellular carcinoma Hep3B cells. J Ethnopharmacol. 2015;166:185–99.

13. Sujana N, Ramanathan S, Vimala V, Sundaram M, Pemaiah B. Antitumour potential of Passiflora incarnata against ehrlich ascites carcinoma. Int J Pharm Pharm Sci. 2012;4(5):10–3.

14. Ma W-D, Zou Y-P, Wang P, Yao X-H, Sun Y, Duan M-H, et al. Chimaphilin induces apoptosis in human breast cancer MCF-7 cells through a ROS-mediated mitochondrial pathway. Food Chem Toxicol. 2014;70(May):1–8.

15. Kapadia GJ, Azuine M a, Tokuda H, Hang E, Mukainaka T, Nishino H, et al. Inhibitory effect of herbal remedies on 12-O-tetradecanoylphorbol-13-acetate-promoted Epstein-Barr virus early antigen activation. Pharmacol Res. 2002;45(3):213–20.

16. Chaparro R. DC, Maldonado Celis ME, Urango M. LA, RojanoI BA. Propiedades quimiopreventivas de Passiflora mollissima (Kunth) L. H. Bailey (curuba larga) contra cáncer colorrectal. Rev Cuba Plantas Med. 2015;20(1):62–74.

17. Perry NB, Albertson NB, Blunt JW, Cole ALJ, Munro MHG, Walker RL. 4-Hydroxy-2-cyclopentenone: An anti-pseudomonas and Cytotoxic Component from Passiflora tetrandra. Planta Med. 1991;57(4):129–31.

18. Gomes SVF, Portugal LA, dos Anjos JP, de Jesus ON, de Oliveira EJ, David JP, et al. Accelerated solvent extraction of phenolic compounds exploiting a Box-Behnken design and quantification of five flavonoids by HPLC-DAD in Passiflora species. Microchem J. 2017;132:28–35.

19. Pereira CAM, Yariwake JH, McCullagh M. Distinction of the C-glycosylflavone isomer pairs orientin/isoorientin and vitexin/isovitexin using HPLC-MS exact mass measurement and in-source CID. Phytochem Anal. 2005;16(5):295–301.

20. Mosmann T. Rapid Colorimetric assay for cellular growth and survival: application to proliferation and cytotoxicity assay. J Immunol Methods. 1983;65(1–2):55–63.

21. Nakayama GR, Caton MC, Nova MP, Parandoosh Z. Assessment of the Alamar Blue assay for cellular growth and viability in vitro. J Immunol Methods. 1997;204(2):205–8.

22. Veras ML, Bezerra MZB, Braz-Filho R, Pessoa ODL, Montenegro RC, Do Ó Pessoa C, et al. Cytotoxic epimeric withaphysalins from leaves of Acnistus arborescens. Planta Med. 2004;70(6):551–5.

23. Geng C-X, Zeng Z-C, Wang J-Y. Docetaxel inhibits SMMC-7721 human hepatocellular carcinoma cells growth and induces apoptosis. World J Gastroenterol. 2003;9(4):696–700.

24. Militão GCG, Dantas INF, Pessoa C, Falcão MJC, Silveira ER, Lima MAS, et al. Induction of apoptosis by pterocarpans from Platymiscium floribundum in HL-60 human leukemia cells. Life Sci. 2006;78(20):2409–17.

25. Mahmoud TS, Marques MR, Pessoa C do Ó, Lotufo LVC, Magalhães HIF, Moraes MO de, et al. In vitro cytotoxic activity of Brazilian Middle West plant extracts. Rev Bras Farmacogn. 2011;21(3):456–64.

26. da Silva TBC, Costa COD, Galvão AFC, Bomfim LM, Rodrigues ACB da C, Mota MCS, et al. Cytotoxic potential of selected medicinal plants in northeast Brazil. BMC Complement Altern Med. 2016;16(1):199.

27. Andrade LN, Lima TC, Amaral RG, Do Ó Pessoa C, De Moraes Filho MO, Soares BM, et al. Evaluation of the cytotoxicity of structurally correlated p-menthane derivatives. Molecules. 2015;20(7):13264–80.

28. Davis BD, Brodbelt JS. Determination of the glycosylation site of flavonoid monoglucosides by metal complexation and tandem mass spectrometry. J Am Soc Mass Spectrom. 2004;15(9):1287–99.

29. Wolfender J, Rodriguez S, Hostettmann K. Liquid chromatography coupled to mass spectrometry and nuclear magnetic resonance spectroscopy for the screening of plant constituents. J Chromatogr A. 1998;794:299–316.

30. Schutz K, Kammerer D, Carle R, Schieber A. Identification and Quantification of Caffeoylquinic Acids and Flavonoids from Artichoke (Cynara. Control. 2004;52:4090–6.

31. Chen X, Zhong D, Jiang H, Gu J. Characterization of some glucuronide conjugates by electrospray ion trap mass spectrometry. Yao Xue Xue Bao. 1998 Nov;33(11):849–54.

32. Pearson CH, Cornish K, McMahan CM, Rath DJ, Whalen M. Natural rubber quantification in sunflower using an automated solvent extractor. Ind Crops Prod. 2010;31(3):469–75.

33. Saha S, Walia S, Kundu A, Sharma K, Paul RK. Optimal extraction and fingerprinting of carotenoids by accelerated solvent extraction and liquid chromatography with tandem mass spectrometry. Food Chem. 2015;177.

34. Batra P, Sharma AK. Anti-cancer potential of flavonoids: recent trends and future perspectives. 3 Biotech. 2013;3(6):439–59.

35. Ganesan K, Xu B. Molecular targets of vitexin and isovitexin in cancer therapy: a critical review. Ann NY Acad Sci. 2017;1401:102–13.

36. Xiao J, Capanoglu E, Jassbi AR, Miron A. Advance on the Flavonoid C-glycosides and Health Benefits. Crit Rev Food Sci Nutr. 2016;56(October):S29–45.

37. Yuan L, Wang J, Xiao H, Xiao C, Wang Y, Liu X. Isoorientin induces apoptosis through mitochondrial dysfunction and inhibition of PI3K/Akt signaling pathway in HepG2 cancer cells. Toxicol Appl Pharmacol. 2012;265(1):83–92.

38. An F, Wang S, Tian Q, Zhu D. Effects of orientin and vitexin from Trollius chinensis on the growth and apoptosis of esophageal cancer EC-109 cells. Oncol Lett. 2015;10(4):2627–33.

39. Patel K, Gadewar M, Tahilyani V, Patel DK. A review on pharmacological and analytical aspects of diosmetin: A concise report. Chin J Integr Med. 2013;19(10):792–800.

40. Méry B, Guy JB, Vallard A, Espenel S, Ardail D, Rodriguez-Lafrasse C, et al. In vitro cell death determination for drug discovery: A landscape review of real issues. J Cell Death. 2017;10.

41. Yuan L, Wei S, Wang J, Liu X. Isoorientin induces apoptosis and autophagy simultaneously by reactive oxygen species (ROS)-Related p53, PI3K/Akt, JNK, and p38 signaling pathways in HepG2 cancer cells. J Agric Food Chem. 2014;62(23):5390–400.

42. Harvey A, Cree I. High-Throughput Screening of Natural Products for Cancer Therapy. Planta Med. 2010;76(11):1080–6.

43. Zong W, Thompson CB. Necrotic death as a cell fate. Genes Dev. 2006;20:1–15.

44. Li YZ, Li CJ, Pinto a V, Pardee a B. Release of mitochondrial cytochrome C in both apoptosis and necrosis induced by beta-lapachone in human carcinoma cells. Mol Med. 1999;5(4):232–9.

